# Intercellular transfer of mitochondria between senescent cells through cytoskeleton-supported intercellular bridges requires mTOR and Cdc42 signalling

**DOI:** 10.1101/2020.11.02.364919

**Authors:** Hannah Walters, Lynne S Cox

## Abstract

Cellular senescence is a state of irreversible cell proliferation arrest induced by various stressors including telomere attrition, DNA damage and oncogene induction. While beneficial as an acute response to stress, accumulation of senescent cells with increasing age is through to contribute adversely to development of cancer and a number of other age-related diseases, including neurodegenerative diseases for which there are currently no effective disease-modifying therapies. Non-cell autonomous effects of senescent cells have been suggested to arise through the SASP, a wide variety of pro-inflammatory cytokines, chemokines and exosomes secreted by senescent cells. Here, we report an additional means of cell communication utilised by senescent cells via large numbers of membrane-bound intercellular bridges - or tunnelling nanotubes (TNTs) - containing the cytoskeletal components actin and tubulin, and which form direct physical connections between cells. We observe the presence of mitochondria in these TNTs, and show organelle transfer through the TNTs to adjacent cells. While transport of individual mitochondria along single TNTs appears unidirectional, we show by differentially labelled co-culture experiments that organelle transfer through TNTs can occur both between different cells within senescent cell populations, and also between senescent and proliferating cells. Using small molecule inhibitors, we demonstrate that senescent cell TNTs are dependent on signalling through the mTOR pathway, which we further show is mediated at least in part through downstream actin-cytoskeleton regulatory factor Cdc42. These findings have significant implications for development of senomodifying therapies, as they highlight the need to account for local direct cell-cell contacts as well as the SASP in order to treat cancer and diseases of ageing in which senescence is a key factor.

## Introduction

Intercellular communication is crucial in regulating cellular function, for example in response to environmental or intracellular stress. Such interplay has historically been thought to be coordinated through secretion of soluble factors including chemokines, cytokines, growth factors and hormones, and their recognition by cell-surface receptors, or through secretion and internalization of extracellular vesicles. However, recent evidence suggests an alternative form of cell-cell communication can be mediated through intercellular membrane connections. Such connections, termed tunnelling nanotubes (TNTs) or intercellular bridges, have been characterized as long, fragile, open-ended and transient protrusions which mediate membrane continuity between connected cells. Since their initial description (Rustom et al. 2004), nanotubes have been observed connecting cells of the same or different cell types, with a particular prevalence detected in immune cells including macrophages, monocytes and NK cells (Dupont et al. 2018). The cargo shuttled within nanotubes includes nutrients, sterols, plasma membrane components, signalling molecules, proteins, RNA species and ions that passively diffuse between connected cells, alongside larger cargo such as whole organelles or protein complexes that require transport by myosin motors (Marzo, Gousset, and Zurzolo 2012). Nanotubes may play important physiological functions, for example in osteoclast differentiation (Takahashi et al. 2013), and have been observed *in vivo*, both in murine myeloid cells in the cornea (Chinnery, Pearlman, and McMenamin 2008) and in human malignant pleural mesothelioma (Lou et al. 2012).

Intercellular bridges can form either by protrusion elongation, where one cell extends filopodia-like protrusions which subsequently connect with a second nearby cell, or by cell dislodgement, where two cells which are initially close move apart, leaving behind a long, thin membrane connection (Dupont et al. 2018). Both result in formation of nanotubes with diameters ranging from 50-1500 nm, spanning tens to hundreds of microns between connected cells. Actin filaments (together with myosin) are present in ‘thin’ TNTs, while ‘thick’ TNTs additionally contain microtubules, raising the possibility that cytoskeletal composition could control TNT cargo transport through size exclusion (Onfelt et al. 2006). Given the universal requirement for F-actin in bridge structures, it is unsurprising that regulators of actin cytoskeletal dynamics including Rac1, Cdc42 and their respective downstream effectors WAVE and WASP are also implicated in bridge formation, as well as the myosin motor protein Myo10 (Hanna et al. 2017). Cdc42 expression, which also regulates filopodia formation, appears important for both bridge elongation and for intercellular cargo transport (Ohno, Hase, and Kimura 2010).

Intercellular communication through TNTs may represent a cellular response to stress, potentially allowing rescue through transfer of functional components from undamaged cells nearby. Consistent with this, both oxidative stress and DNA damage are associated with increased TNT formation (Domhan et al. 2011), in a manner dependent on p53 (Wang et al. 2011). Stressed cells appear to ‘reach out’ to unstressed cells (Wang et al. 2011) and transfer of mitochondria from healthy neighbouring cells to stressed cells via TNTs has been speculated to serve as a rescue mechanism for stress tolerance. For example, intercellular mitochondrial transport allows survival of cancer cells experiencing loss of mitochondrial functionality (Wang and Gerdes 2015). As well as providing rescue from metabolic failure or mitochondrial dysfunction, mitochondrial transfer via TNTs has even been reported to result in cellular reprogramming (Koyanagi et al. 2005). Nanotube formation may also be involved in induction of apoptosis, as the death signal Fas ligand has been noted to be shuttled via TNTs in T lymphocytes to induce cell death in target cells (Arkwright et al. 2010).

Stress is therefore associated with TNT formation, but cellular stress is also a known driver of cell senescence (reviewed by (de Magalhães and Passos 2018). It is hence noteworthy that intercellular membrane connections have been observed to increase upon induction of cell senescence, with transfer of cytoplasmic proteins preferentially from senescent cells to natural killer (NK) cells in co-culture (Biran et al. 2015), resulting in increased NK cell activation and cytotoxicity. Proteomic analysis of transferred cargo showed transfer of proteins implicated in actin re-organization. In particular, Cdc42 was reported to be substantially upregulated and highly active in senescent versus proliferating fibroblasts, while its inhibition substantially reduced protein transfer and NK cytotoxicity (Biran et al. 2015).

Here, we set out to investigate the composition and role of intercellular bridges in cell senescence, and whether the bridges are capable of supporting transfer of organelles between cells. We report a high prevalence of membrane-bound TNTs formed by senescent cells, containing both actin and tubulin. We further show that mitochondria can be transferred through these bridges and that mTOR signalling and Cdc42-mediated actin organisation pathways are critical for organelle transfer through tunnelling nanotubes in senescent cells. These findings highlight a potential target for new therapies directed against senescent cells.

## Materials and Methods

### Cell culture

HF043 neonatal foreskin fibroblasts (Dundee CELL products) were verified to be primary diploid human fibroblasts, uncontaminated with any known lab cell line, by short tandem repeat analysis (Porton Down, UK). IMR90 ER:Ras fibroblasts (16 week female foetal lung fibroblasts) were a gift from Prof. Peter Adams (University of Glasgow, UK, and Sanford Burnham Prebys Medical Discovery Institute, La Jolla, USA). Cells were cultured in DMEM (D5796, Gibco) supplemented with 10% heat-inactivated FCS (Gibco) for HF043, or 1 mM sodium pyruvate (Gibco) and 20% FCS for IMR90 ER:ras fibroblasts. (Media was not supplemented with antibiotics). Cells were subcultured once they reached ~70% confluence as assessed using a digital EVOS microscope (Thermo Fisher), by washing in PBS (Sigma), 3-5 minute incubation with Tryple Express Trypsin (Thermo Fisher), and dilution and gentle trituration in complete media. Cell viability and number were assessed using a T4 Cellometer (Nexelcom), from which population doublings were subsequently calculated as PD = (log[number harvested/number seeded])/log(2). Cells were seeded at 4-8×10^3^ cells/cm^2^ in filter-capped flasks or multi-well plates (Greiner) and incubated with complete media in a humidified incubator at 37oC in 5% CO_2_ and 20% O_2_. Cells were regularly inspected by phase contrast microscopy for cell health, and tested for mycoplasma contamination by PCR according to the method of Uphoff and Drexler (Uphoff and Drexler 2002, 2004).

### Drug treatments

All drugs used were reconstituted and stored as directed by the supplier (Etoposide, 4-OHT, Casin - all Sigma-Aldrich; AZD8055 - Selleckchem). For routine drug treatment, cells were seeded in complete media and allowed to bed down overnight, before media was aspirated and replaced with drug-supplemented complete media. For drug treatment in co-culture assays, cells were seeded directly into drug-supplemented complete media. Optimum concentrations and dosing periods were determined empirically from literature-informed pilot experiments (not shown).

### Induction of senescence

Replicative senescence (RS): the primary human fibroblast line HF043 was grown in continuous culture until replicative exhaustion. Cells were determined to be replicatively senescent when populations fulfilled each of the following criteria: failure to increase in cell number within >2 weeks, a cumulative population doubling number of >85, and positive SA-β-gal staining according to manufacturer’s instructions (Cell Signaling Technology #9860S).

DNA damage-induced senescence (DDIS): cells were treated with 20 μM etoposide for 7 days and verified as senescent by SA-β-gal staining and morphological assessment, as well as failure to re-proliferate. Cells at low CPD (<50) were always used for etoposide treatment to avoid confounding effects of replicative senescence.

Oncogene-induced senescence (OIS): IMR90 ER:Ras cells were incubated with 1 μM 4-OHT for 7 days and assessed for senescence induction by SA-β-gal staining, morphological assessment and failure to re-proliferate.

### Fluorescence staining

For live imaging of mitochondria, cells were incubated for 30 minutes with Mitotracker Green FM or Mitotracker Red (1:1000 v/v dilution of 1 mM stock) according to manufacturer’s instructions (Invitrogen Molecular Probes), in complete media at 37°C in the dark. Media was replaced before imaging to avoid fluorescent flare. Alternatively, cells were incubated overnight with GFP-BacMam probe for mitochondria (GFP expressed fused to the leader sequence of E1 α pyruvate dehydrogenase) according to manufacturer’s instructions (CellLight, Invitrogen). For co-culture experiments, cells were stained with appropriate label *in situ*, washed 3x in PBS, harvested by Trypsin treatment (as above) before re-seeding at 1:1 ratios. To assess potential confounding dye leakage, conditioned media was harvested at 24h from stained cells and incubated for ≥24h with control unstained cells.

For analysis of fixed samples, mitochondria were stained for mitochondrial-specific TFAM and analysed by immunofluorescence. Briefly, cells were washed in PBS, fixed in 3.7% formaldehyde (10 minutes, RT), washed twice in PBS, blocked in 5% donkey serum (Dako) in PBS and incubated with primary antibody (α-TFAM (Mouse) Ab, Abnova B01P) diluted 1:200 in PBS containing 0.3% Triton X-100 v/v and 1% BSA, w/v, at 4°C overnight in a humidified chamber. Cells were then washed twice in PBS and incubated with secondary antibody (Alexafluor 488 α-mouse IgG (Donkey), Invitrogen A-2102 2 hours, RT in the dark, 1:500 in dilution buffer as for primary antibody). Cells were washed twice in PBS and DNA counter-stained before imaging.

For F-actin staining, cells were washed in PBS then fixed in 3.7.% formaldehyde in PBS (v/v, 10 minutes RT) before washing in PBS, then incubated for 40 minutes with FITC-phalloidin (6.6 nM in PBS, Molecular Probes, Thermo Fisher). Cells were then washed in PBS and imaged. For tubulin staining, cells were incubated with Tubulin Tracker Green diluted in complete media according to manufacturer’s instructions (30 minutes, 37°C, Thermo Fisher), media was replaced, and cells were imaged live.

Plasma membranes were labelled by incubating cells with 1:250 wheat germ agglutinin (WGA, v/v) conjugated with the fluorophore FITC or rhodamine (Vector Labs), either live in complete media (immediately after staining to avoid WGA-induced cytotoxicity) or after 3.7% formaldehyde fixation in PBS (without permeabilization). Our optimisation studies comparing live imaging with fixed cells suggested that while fixation improved the sharpness of cell staining, it did so without disrupting the intercellular bridge structures (data not shown).

DNA was stained using NucBlue Live (Hoechst 33342) according to manufacturer’s instructions (20 minutes, 1 drop/ml, RT, Thermo Fisher) or using Hoechst 33342 within mounting dye (VECTACHSIELD, Vector Labs) for fixed cells, with 1 drop of mounting medium used for coverslip mounting.

### Microscopy

Phase contrast microscopy was performed using a digital EVOS Core microscope (Thermo Fisher), with scale bars added to images using a graticule. Phase contrast microscopy was used for routine assessment of cells to determine % confluence before subculturing, as well as for morphological analysis (all images taken using a 20X lens). Fluorescence microscopy was performed using a ZOE fluorescent cell imager (BioRad), with scale bars added automatically. For time-lapse imaging, the field of view was locked and images taken at specified intervals (see specific Figures) according to a stopwatch. Within individual experiments, optical gain was fixed at the outset of image acquisition to ensure image-to-image comparability. Between experiments, brightness and (where stated) sharpness were normalised across different samples within Powerpoint. Where necessary FUJI software was used to enhance colour discrimination (Figure 3).

### Statistical Analysis

The GraphPad Prism 8 statistical analysis package was used to perform statistical tests. For comparisons between >2 samples, ANOVAs (analysis of variance) were performed with Tukey tests to compare the means of each sample. All graphs show mean values with standard deviations (as error bars), calculated from n≥3 independent experiments or from technical triplicates from representative experiments of n≥3 unless otherwise stated.

## Results

### Intercellular bridges connect senescent cells

To examine whether intercellular bridges are associated with cell senescence, we induced replicative senescence (RS) in primary human skin fibroblasts (HF043) by longitudinal continuous cell culture (serial passaging) or induced DNA-damage induced senescence (DDIS) by treating cells with etoposide for 7 days (see Methods). In parallel, we treated IMR90 ER:RAS lung fibroblasts with tamoxifen (4-OHT) to trigger oncogene-induced senescence (OIS), and compared these with proliferating control cells. As expected, the proliferating cells showed characteristic spindle-like morphology, relatively small size and were mononuclear (Figure 1A), while the senescent cells by contrast were greatly enlarged, contain granular inclusions, were often multinucleate with prominent nucleoli, and cell margins were poorly defined under phase contrast optics (Figure 1B).

**Figure 1.**
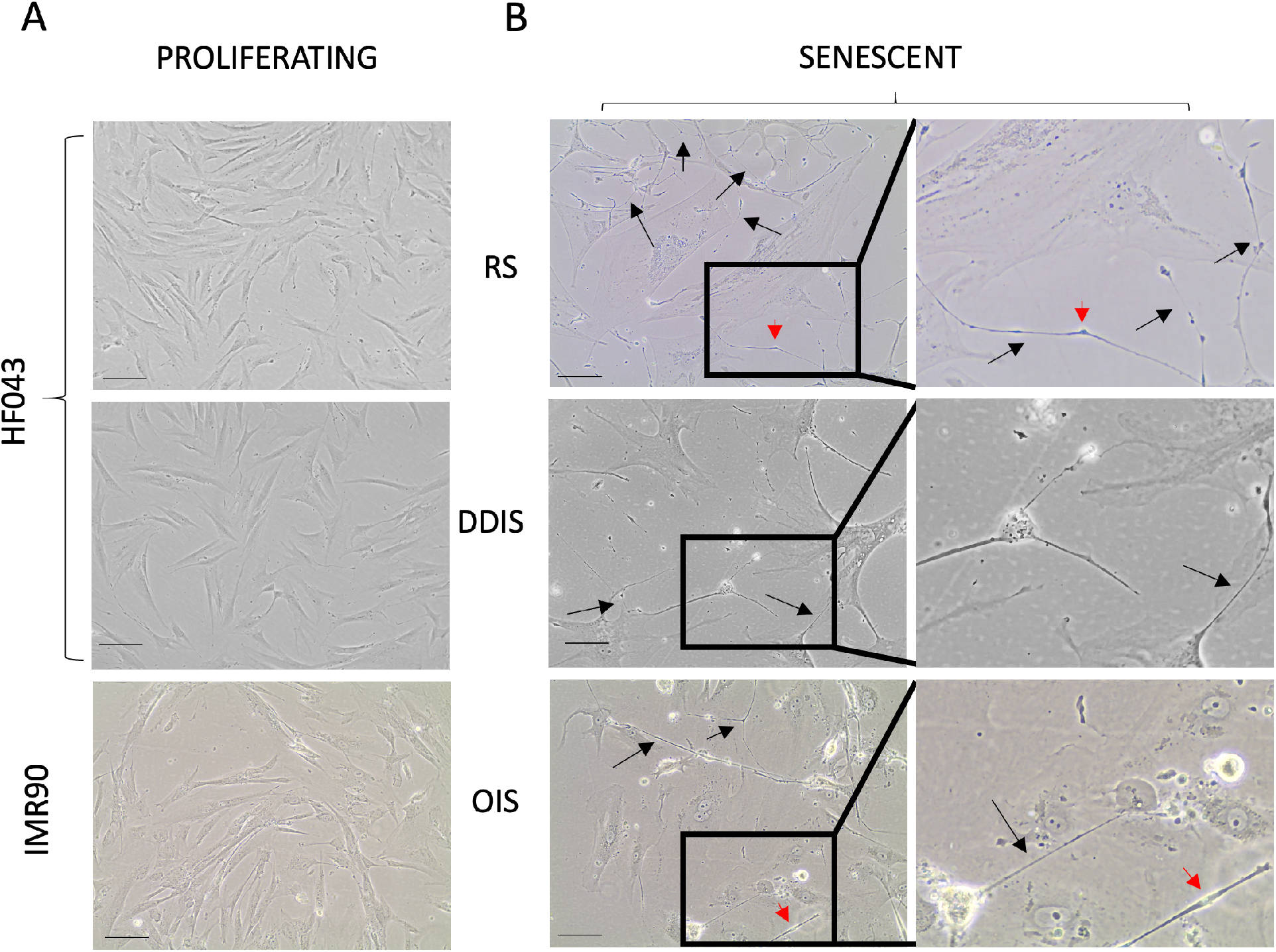
Senescent cells are connected by thin intercellular contacts *in vitro*. (A) Phase contrast images of proliferating HF043 fibroblasts at low cumulative population doubling (CPD<40), and vehicle control proliferating IMR90 ER:ras fibroblasts; (B) HF043 fibroblasts that have undergone replicative senescence (CPD >90) or DNA damage-induced senescence (DDIS, following 7 day 20 μM etoposide treatment), together with oncogene-induced senescent (7 day 4-OHT) ER:ras fibroblasts. Black arrows indicate examples of intercellular bridges, red arrowheads indicate bulky protrusions within the bridges. Representative data shown of n>3 experiments. Scale bar 100 μm.

We observed by phase contrast microscopy a substantial number of long, thin structures of considerable length (tens to hundreds of microns) directly connecting nearby cells within senescent populations of both skin (HF043) and lung (IMR90) fibroblasts, irrespective of mode of senescence induction (Figure 1B, bridges indicated by black arrowheads). The bridge-like structures that we observe in the senescent cell populations were infrequent within proliferating control populations (Figure 1A), in agreement with previous data suggesting nanotube formation occurs in response to cellular stress (Biran et al. 2015) since senescence is a stress response. We further noted a number of swellings at various points along the nanotubes (red arrowheads, Figure 1B). These appear similar to protrusions previously described as ‘gondolas’ within nanotubes, that are thought to be associated with transport of large cargo (Veranic et al. 2008) including organelles such as mitochondria.

### Bridges are membrane-bound and contain actin and tubulin

To determine whether the observed senescent cell bridges are membrane-bound, we stained cells with FITC-conjugated lectin wheat germ agglutinin (WGA) to assess the presence of the sugar O-Glc-NAc, which is prevalent on mammalian membranes; DNA was counterstained with NucBlue-Live. From Figure 2A, multiple membrane-bound (WGA-positive) protrusions can be seen to extend between the senescent cells towards their neighbours, in such a way that the cells appear to form a ‘network’ of interconnected cells rather than discrete cells. The diameter of such connections varied from ultrafine bridges (Figure 2B) reminiscent of ‘thin’ TNTs, and larger diameter bridges (Figure 2C) that appear consistent with previously described ‘thick’ TNTs (Onfelt et al. 2006).

**Figure 2.**
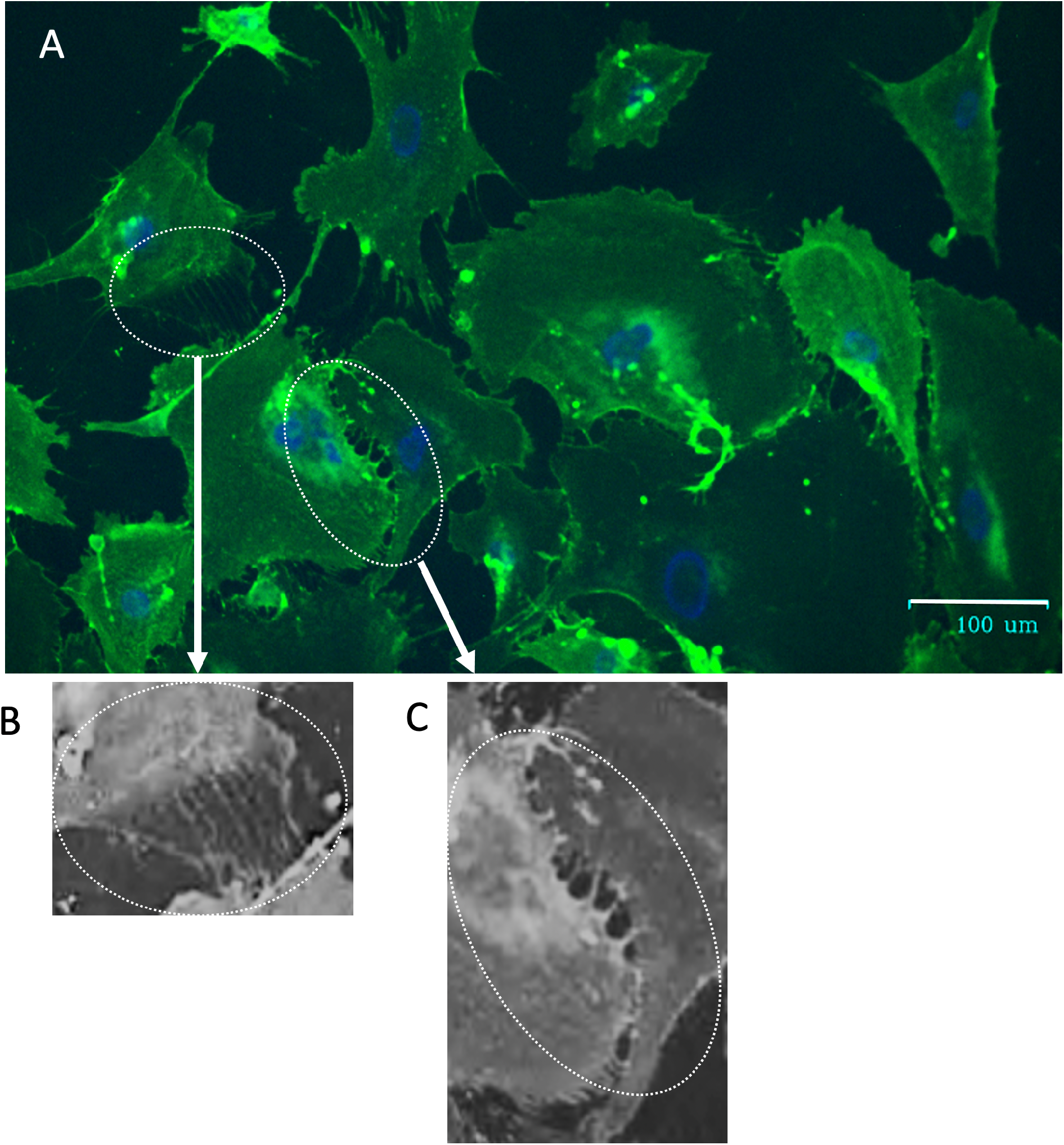
Intercellular bridges are membrane-bound and occur at high frequency between senescent cells. (A) HF043 fibroblasts cultured to replicative senescence and live imaged with FITC-WGA to highlight membrane-associated O-Glc-NAc and NucBlue Live for DNA. (B, C) Magnified images of thin bridges (B), and larger diameter bridges (C). Images in B and C have been recoloured in grayscale and sharpness-enhanced to show the bridges more clearly.

To determine whether these structures represent actin-stabilised tunnelling nanotubes (TNTs), formaldehyde-fixed cells were co-stained with rhodamine-WGA (for membranes) and FITC-phalloidin for actin, and counterstained with NucBlue-Live for DNA (Figure 3A), or live-imaged with Tubulin Tracker Green to identify polymerized tubulin (Figure 3B). Fluorescence microscopy demonstrated that the intercellular bridges contain actin (Figure 3A) and tubulin (Figure 3B); in both cases, the bridges stained positively with rhodamine-WGA, demonstrating the presence of O-Glc-NAc, orthogonally confirming with a different dye our observations in Figure 2. These data suggest that senescent fibroblasts form intercellular nanotubes that contain actin and/or tubulin.

**Figure 3.**
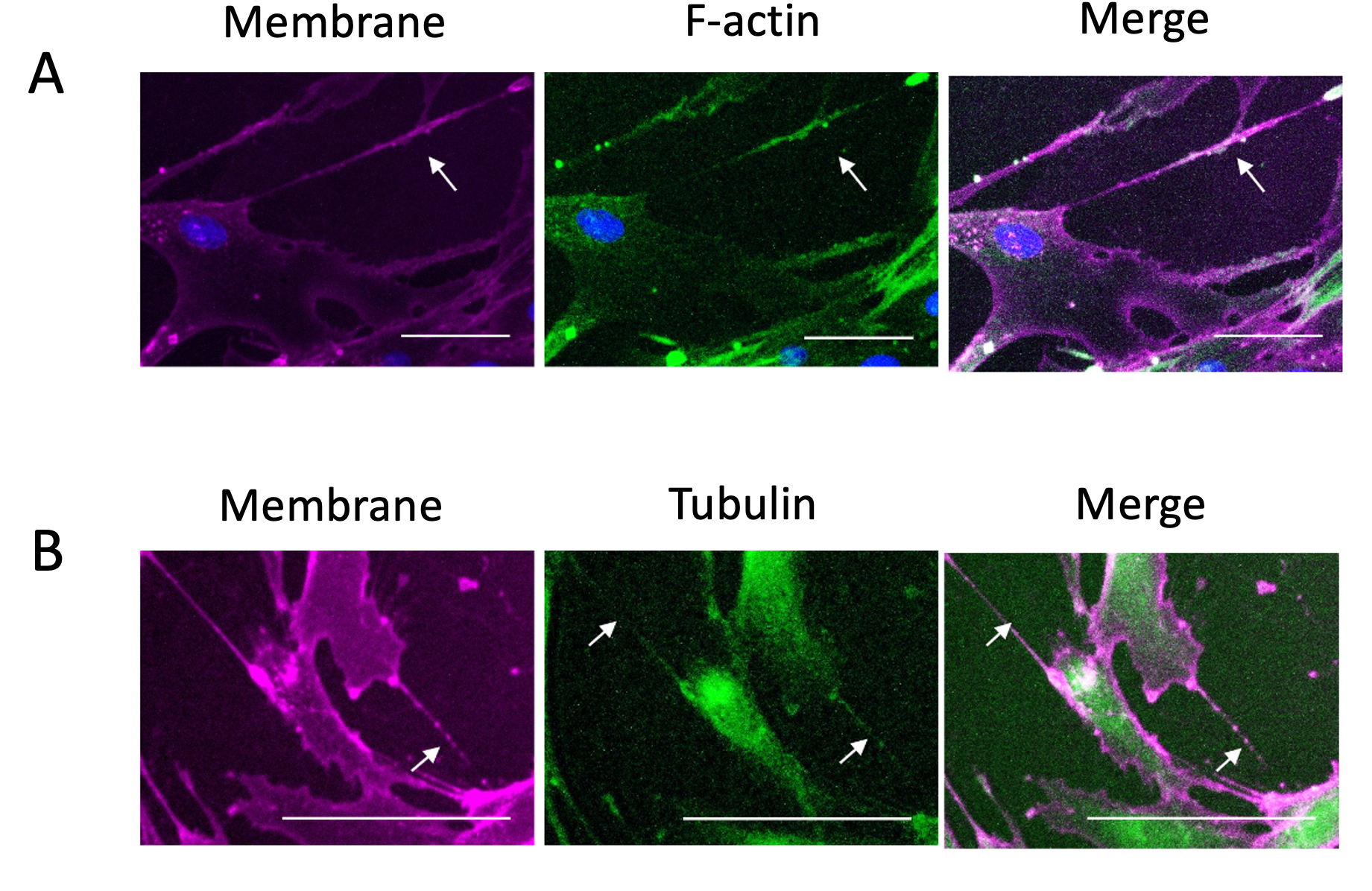
Intercellular contacts between senescent cells contain actin and tubulin. Replicatively senescent HF043 fibroblasts (CPD >90) were (A) fixed and stained with rhodamine-WGA (membrane, purple), FITC-phalloidin (F-actin, green) and NucBlue Live (DNA, blue) or (B) live imaged with rhodamine-WGA (membrane, purple) and Tubulin Tracker Green (microtubules, green) prior to analysis by fluorescence microscopy. N=3, representative images shown. Arrows indicate examples of intercellular bridges. Images have been false coloured from the original red/green in FUJI to improve dye discrimination. Scale bar 100 μm.

### Mitochondria are present in senescent cell TNTs

As signalling hubs and the major source of oxidative stress, mitochondria play an important role in senescence (Correia-Melo et al. 2016). Senescent cells exhibit dramatic increases in mitochondrial load (Walters, Deneka-Hannemann, and Cox 2016), possibly to compensate for mitochondrial dysfunction, and the mitochondrial network becomes increasingly reticular through resistance to mitochondrial fission and mitophagy (Korolchuk et al. 2017).

To ask whether the senescent cell TNTs are capable of transporting mitochondria, we labelled mitochondria in senescent fibroblasts (RS and DDIS in HF043, and OIS in IMR90, as above) with fluorescent dye. To avoid labelling bias, we compared a variety of mitochondrial dyes including the live stains Mitotracker Green, Mitotracker Red, and CellLight Mitochondria GFP BacMam, where GFP is localised to mitochondria through fusion of GFP with the leader sequence from E1 alpha pyruvate dehydrogenase. We also examined staining patterns of the mitochondrial transcription factor TFAM in fixed samples. In all cases, cell membranes were stained with fluorescent WGA (dye colour indicated in labels of Figure 4).

**Figure 4.**
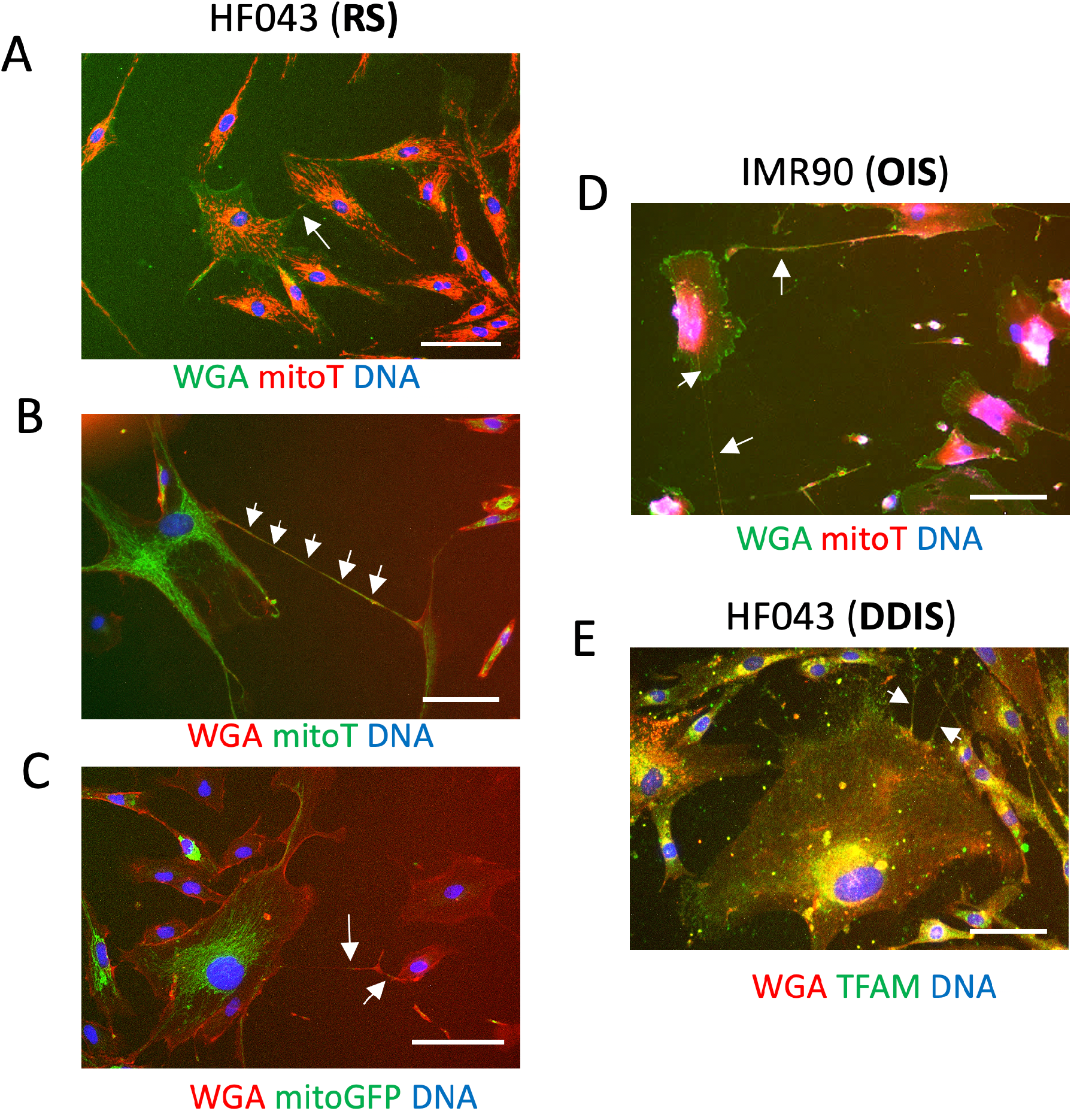
Mitochondria are present within senescent intercellular contacts. (A-C) Replicatively senescent HF043 fibroblasts were stained with (A) fluorescein-WGA and Mitotracker Red (mitoT), rhodamine-WGA and Mitotracker Green (mitoT), (C) rhodamine-WGA and CellLight Mitochondria-GFP BacMam (mitoGFP). (D) IMR90 ER:ras cells induced to undergo oncogene-induced senescence (OIS) by 7d treatment with 4OHT were stained with fluorescein-WGA and Mitotracker Red (mitoT). Cells in A-D were imaged live. (E) HF043 fibroblasts treated for 7d with etoposide to drive DNA damage-induced senescence (DDIS) were fixed and stained with anti-TFAM primary antibody with Alexafluor 488 secondary antibody and rhodamine-WGA. Arrows indicate mitochondria within contacts. n>3, DNA stained with NucBlue Live. Scale bar 100 μm.

As shown in Figure 4, we observed small foci (Figure 4A, C) or even reticular networks (Figure 4B, D) positive for mitochondrial staining in a substantial proportion of intercellular bridges, irrespective of the mitochondrial stain used, the mode of senescence induction, or whether cells were imaged live or fixed, showing the presence of mitochondria within TNTs of senescent cells.

### Senescent cell tunnelling nanotubes support intercellular mitochondrial transport

The mitochondria present in intercellular bridges may arise simply through a chance peripheral distribution at the time and subcellular location of bridge formation; alternatively, they could represent cargo being transported through the bridges, as previously suggested (Wang and Gerdes 2015). Hence to assess whether the observed mitochondria were static or mobile within these intercellular bridges, time-lapse fluorescence microscopy of replicatively senescent HF043 fibroblasts stained with Mitotracker Red was conducted.

Mitochondrial motility was observed within minutes of staining and mitochondria were seen to track along intercellular bridges, as seen in representative still images of time lapse microscopy (Figure 5). All movement detected for individual mitochondrial puncta was unidirectional, with a relatively constant speed of ~1μm/minute within the observation period; the total distance travelled by the mitochondria in the example shown in Figure 5 was ~60 μm within 70 minutes.

**Figure 5.**
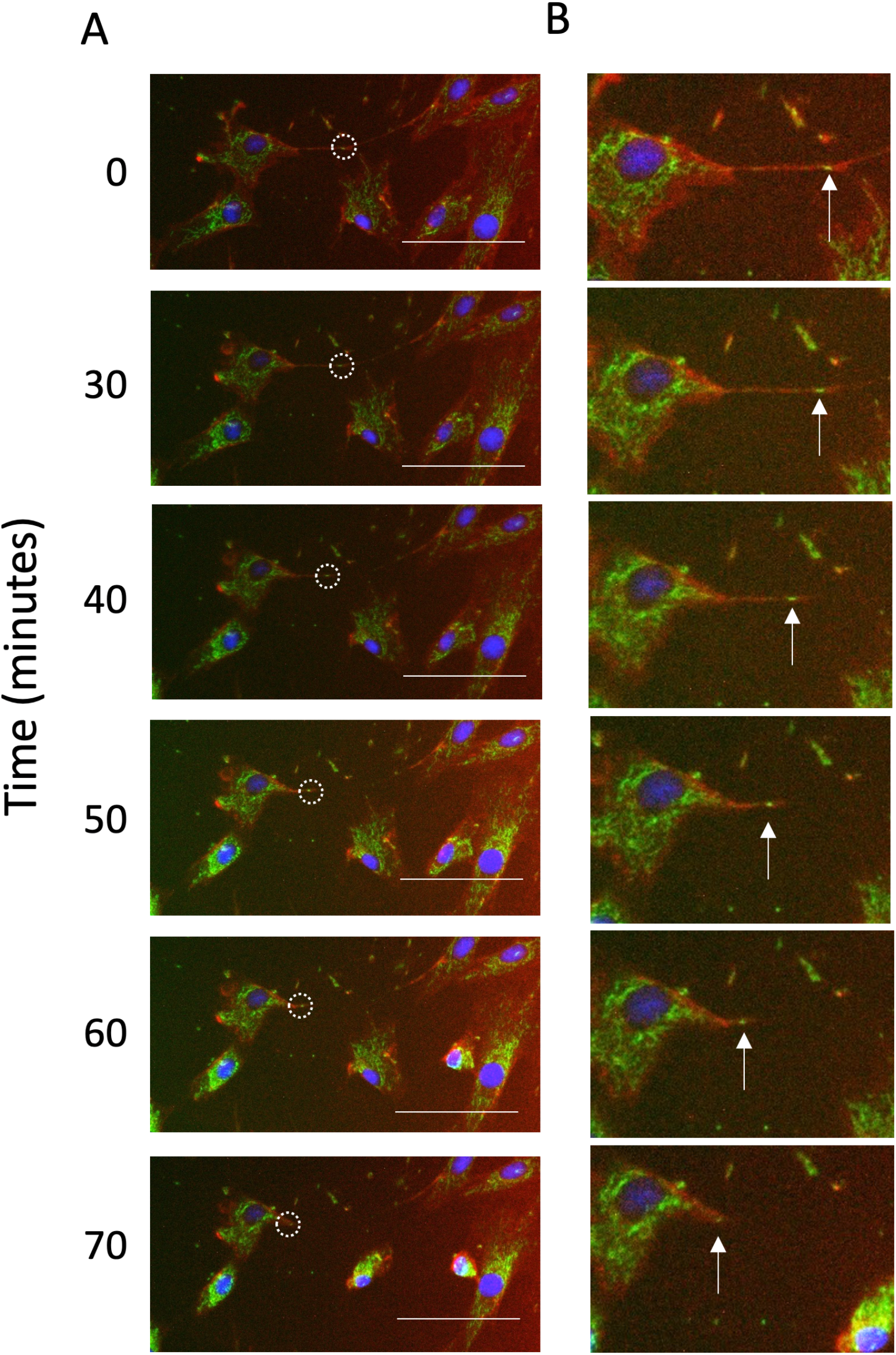
Mitochondria are motile within senescent intercellular bridges. Replicatively senescent HF043 fibroblasts were stained with Mitotracker Green, rhodamine-WGA and NucBlue Live (for DNA) and imaged live using time-lapse fluorescence microscopy, with time points following the start of the observation period indicated in minutes. The white dotted circle highlights a Mitotracker Green positive punctum within a TNT. Representative images shown of n=3 experiments. Scale bar 100 μm. (B) Magnification of TNT with position of moving Mitotracker positive puncta indicated by arrow.

These observations suggest that mitochondria are able to move within senescent intercellular bridges over the timescale of minutes to hours. The observation of unidirectional movement also suggests that mitochondrial motility may be regulated, rather than occurring by passive diffusion, possibly involving motor proteins acting on the underlying cytoskeletal framework of actin and/or tubulin.

### Co-culture demonstrates transfer of mitochondria between cells

To investigate whether this putative mitochondrial transport resulted in transfer of mitochondria between cells, we next conducted a co-culture assay where separate populations of cells were stained with Mitotracker Green and Mitotracker Red respectively, then thoroughly washed before harvesting and co-seeding at a 1:1 ratio (schematic shown in Figure 6A, with images of co-seeded cells immediately after plating in Figure 6B).

**Figure 6.**
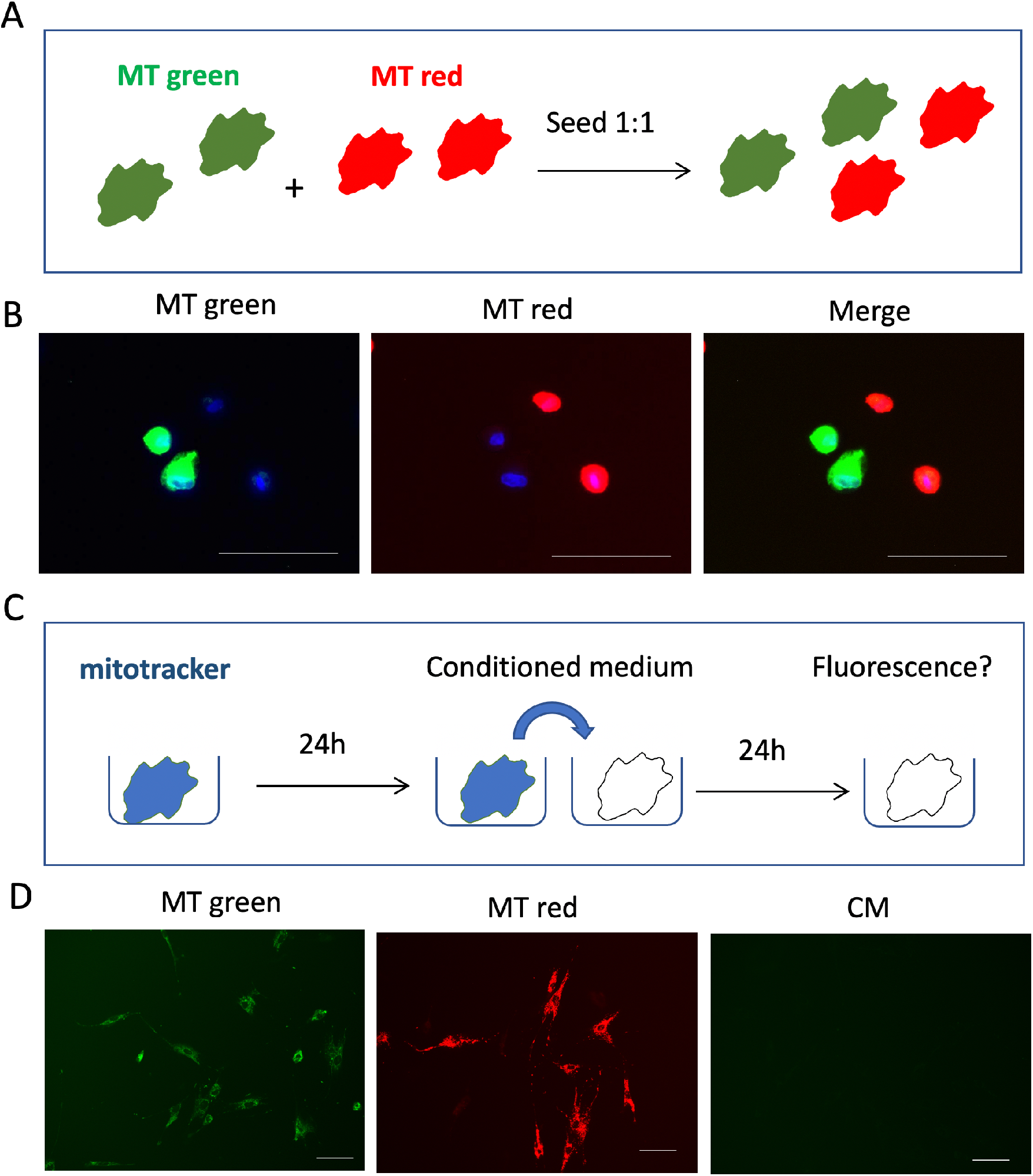
Co-culture assay for analysis of intercellular mitochondrial transfer. (A) Schematic of co-culture set-up. HF043 fibroblasts were stained with Mitotracker Green (MT Green) or Mitotracker Red (MT Red), washed thoroughly and then seeded into co-culture at 1:1 ratio. (B) Cells were imaged immediately after co-plating (proliferating cells shown). (C) Schematic of assay for dye leakage: cells were stained for 30 min with mitotracker green or red, media replaced and incubated for 24h to generation conditioned medium (CM). This CM was then harvested and incubated with unstained cells for 24h prior to fluorescence microscopy. (D) Representative images of proliferating cells stained with Mitotracker green or Mitotracker red, or unstained cells incubated for 24 hours with conditioned media (CM) harvested from stained cells. n>3. Scale bar 100 μm.

To exclude the possibility of mitotracker dye carry-over between different cell populations in co-culture experiments through dye leakage into the medium, conditioned media (CM) controls were included in each experiment whereby media was harvested after 24 hours of exposure to stained cells (either mitotracker green or red, shown simply as ‘mitotracker’ in schematic Figure 6C), followed by incubation with unstained cells for 24 hours with inspection for any fluorescent signal by microscopy; no evidence of dye leakage was detected (Figure 6D).

Having optimised and verified labelling and co-culture conditions as shown in Figure 6, we then assessed whether mitochondria were transferred between differentially labelled cell populations. One population of senescent HF043 cells was labelled with Mitotracker red (MT red) and another with Mitotracker green (MT green), harvested, co-seeded at 1:1 then imaged after 24 hours. Any transfer of mitochondria between cells of different populations might be visualised as discrete puncta of the opposite dye colour, as shown in the schematic (Figure 7A). Indeed, such puncta were detected in a number of co-cultured senescent cells (white arrows, Figure 7B), irrespective of the Mitotracker dye used. In addition, we also observed what appear to be mitochondrial reticular networks extending through TNTs between cells over considerable distances (>100 μm, white arrows Figure 7C), with large senescent ‘donor’ cells also appearing to accept mitochondria from oppositely-stained cell populations (yellow arrow, Figure 7C). We further observed instances where TNS were stained both with red and green, suggesting formation by fusions of TNTs extruded from separate cells (blue arrow, Figure 7C). Mitochondria were also transferred through TNTs from labelled to unlabelled cells (Figure 7D, E).

**Figure 7.**
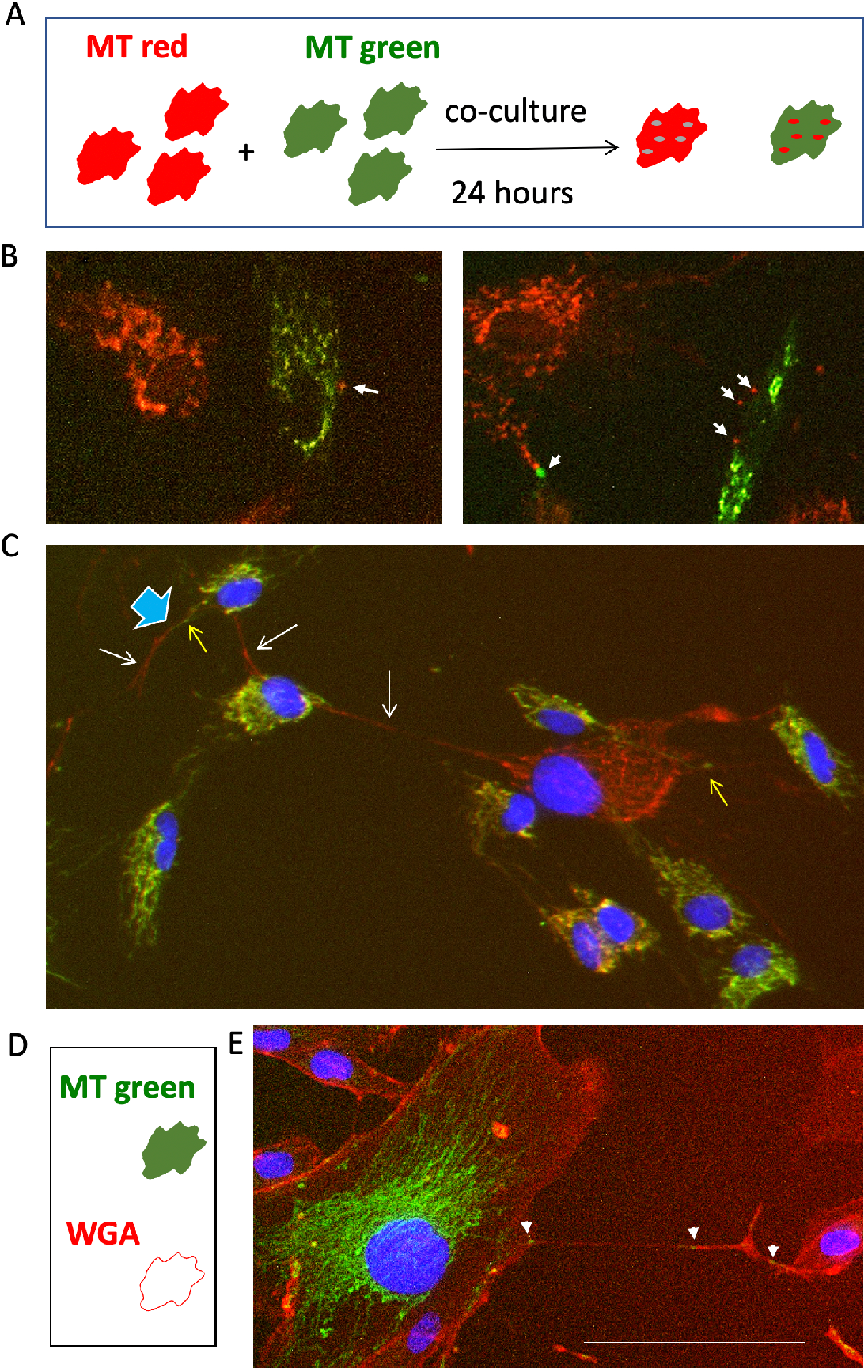
Mitochondrial transfer between co-cultured senescent cells. (A) Schematic of coculture experiment. One senescent cell population is labelled by incubation with mitotracker (MT) red and another with MT green for 30 mins. Following washing and harvesting, the differently labelled cells are seeded at a 1:1 ratio and co-cultured for 24h prior to florescence microscopy analysis. (B) Examples of mitochondrial puncta arising through transfer between cocultured cells (white arrow heads). (C) Example of TNTs that appear to contain reticular mitochondria. White arrows indicate TNTs arising from cell labelled with mitotracker red, yellow arrow those arising from cells labelled with mitotracker green. Blue arrow head indicates TNT that appears to be formed by fusion of a bridge between two differently labelled cells. (D) Schematic showing co-culture between senescent cells labelled with mitotracker (MT) green and unlabelled cells. (Rhodamine WGA was used subsequently to highlight cell membrane)s. (E) Representative image of co-culture as in (D). White arrow heads indicate presence of green mitochondrial puncta being transferred to a cell without mitochondrial labelling. Scale bar 100μm.

### Mitochondria pass from senescent cells to proliferating cells and vice versa

Intercellular mitochondrial transfer has been hypothesized as a rescue mechanism for stressed cells, including cancer cells that have lost mitochondrial functionality (Wang and Gerdes 2015). To determine whether, like stressed cancer cells, senescent cells can accept mitochondria from healthy proliferating cells, we conducted co-culture experiments between populations of proliferating and senescent cells, with ‘red’ stained proliferating cells co-cultured with ‘green’ stained proliferating cells (PRO-PRO), ‘red’ senescent with ‘green’ senescent cells (SEN-SEN) or proliferating and senescent (PRO-SEN) co-cultures (using both colour combinations to eliminate any effect of dye bias) - dye colours are indicated in labels of the Figure 8. Nuclei were counter-stained using the live dye NucBlue Live prior to imaging.

**Figure 8.**
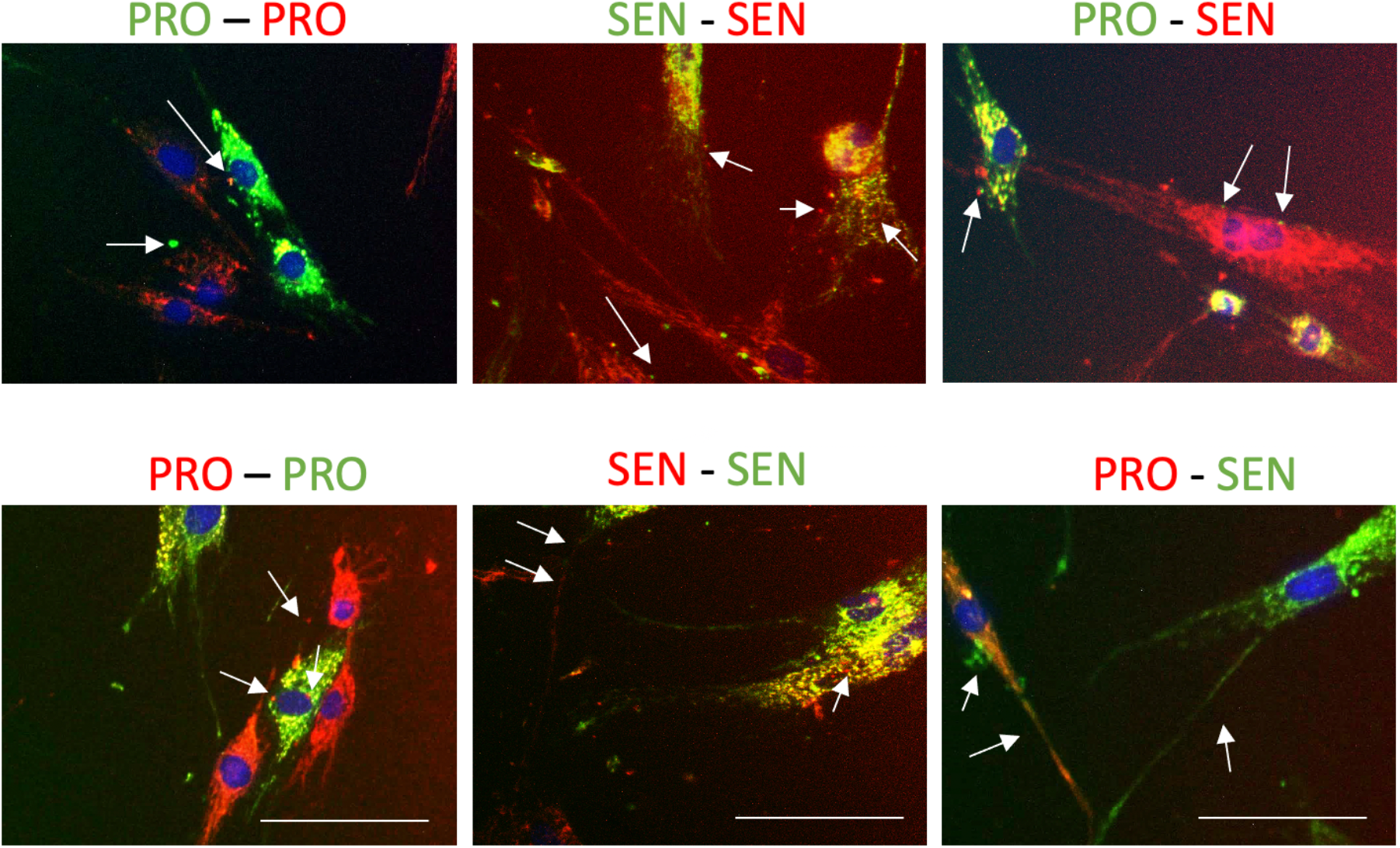
Intercellular mitochondrial transfer occurs between both proliferating and senescent cells. Co-cultures of proliferating (PRO) or replicatively senescent (SEN) HF043 fibroblasts pre-stained with Mitotracker Green or Red were (set up as in Figure 6A) and imaged following 24 hours co-incubation. Arrows indicate intercellular bridges or puncta of transferred mitochondria. N=3, DNA was stained with NucBlue Live immediately prior to analysis. Scale bar 100 μm.

After 24 hours of co-culture, we observed puncta of oppositely stained mitochondria within both proliferating and senescent fibroblasts, regardless of which population was stained with Mitotracker green or Mitotracker red (Figure 8), as well as mitochondria-loaded bridges spanning alternately stained ‘red’ and ‘green’ cells, suggesting that mitochondria have been transferred between different cells, with proliferating and senescent cell populations able to act both as mitochondrial donors and acceptors. Taken together, our results suggest that tubulin- and actin-based intercellular bridges may constitute an important mechanism of mitochondrial transfer and thus intercellular communication for senescent cells, and between senescent and proliferating cells within a tissue.

### Direct intercellular communication in senescence is regulated by mTOR and Cdc42 signalling

Intercellular bridge structures between cancer cells have been reported to be regulated by mTOR signalling (Lou et al. 2012), while actin regulatory factors have been implicated in senescent cell TNTs (Biran et al. 2015). We therefore tested whether the intercellular bridges observed here are regulated by Cdc42 or mTOR signalling, using the pharmacological inhibitors CASIN and AZD8055 to inhibit Cdc42 and mTOR respectively, in both senescent and proliferating cell populations. As an ATP-competitive mTOR inhibitor, AZD8055 impacts not only on pathways regulated by mTORC1 (such as translation and autophagy) but also on those controlled by mTORC2, including actin polymerisation (e.g. (Walters, Deneka-Hannemann, and Cox 2016)).

Populations of proliferating or senescent HF043 fibroblasts which had undergone either replicative senescence (RS) or DNA-damage induced senescence (DDIS) through 7 day treatment with 20 μM etoposide, were exposed to CDC42 inhibitor CASIN at 2 μM, pan-mTOR inhibitor AZD8055 (Walters, Deneka-Hannemann, and Cox 2016) at 70 nM, or DMSO (vehicle control) for 24 hours prior to fixation with formaldehyde. Cells were then stained with fluorescein-WGA for cell membranes and NucBlue live for nuclear DNA. The number of intercellular bridges connected at both ends was counted manually, and number per cell determined by normalising against nuclear number (Figure 9A).

**Fig 9.**
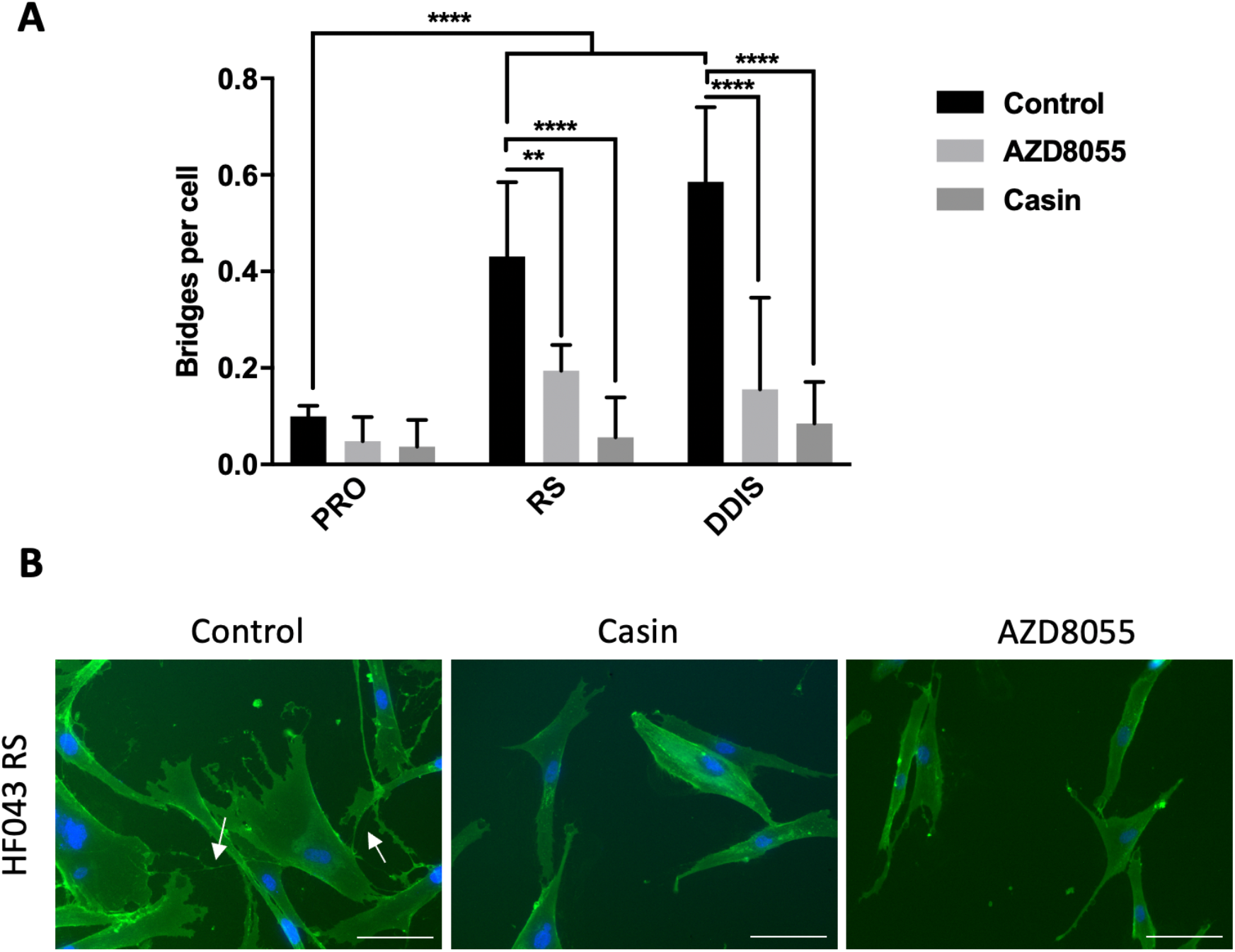
Cdc42 and mTOR signalling are required for formation of intercellular bridges in senescent cells. (A) HF043 fibroblasts that had undergone replicative senescence (RS, cumulative population doubling ≥90), DNA damage-induced senescence (DDIs, following 7 day treatment with 20 μM etoposide), and proliferating control cell populations at low cumulative population doubling (CPD) were treated with the mTOR inhibitor AZD8055 (70 nM) or the Cdc42 inhibitor CASIN (2μM) for 24 hours before fixation and staining with fluorescein-WGA (for membranes) and NucBlue Live (for DNA). Intercellular bridges were manually quantified from >50 cells per replicate (n=3). ** p<0.05, *** p<0.001. (B) Representative images of control and drug-treated replicatively senescent fibroblasts. Arrows indicate intercellular bridges (TNTs) Scale bar = 100 μm.

We observed significantly more intercellular bridges within senescent (RS and DDIS) compared with proliferating control populations (PRO, Figure 9A), with ~0.5 bridges per replicatively senescent (RS) cell. While a substantial proportion of cells exhibited several connections, others had none. Notably disruption of CDC42 signalling upon CASIN treatment, or inhibition of mTOR signalling with AZD8055, each significantly reduced the number of bridges between senescent cells, both in replicatively senescent populations or those induced by DNA damage (Figure 9A); representative fluorescence images of treated RS cells are shown in Figure 9B. We further observed that the reduction in bridge number occurred in a dose-dependent manner (supplementary Figure S1) strongly suggesting that mTOR and CDC42 are centrally important in TNT formation and or stability.

## Discussion

Entry into senescence is accompanied by development of a number of distinctive phenotypes including epigenetic alterations, lysosomal and mitochondrial dysfunction, pro-inflammatory secretion through the SASP, and morphological alterations including hypertrophy and changes to actin organization networks. Furthermore, senescent cells are highly communicative, alerting immune cells to their presence for clearance (Ovadya et al. 2018), and regulating the proliferative capacity of surrounding healthy or cancerous cells positively and negatively (Gonzalez-Meljem et al. 2018, Nelson et al. 2012). While this interplay has largely been attributed to secretion of exosomes and SASP factors, we show a high prevalence of intercellular bridges between senescent cells that may also be important in mediating these effects.

### Supercellularity in senescence?

While intercellular nanotubes appear to constitute a highly localized form of communication, it is important to note that these bridges often do not just connect two cells, but instead can form networks extending direct cell communication over larger distances. Indeed, we have observed cellular networks that appear almost syncytial in nature, with multiple connections between multiple cells (e.g. Figure 2). Hence through allowing transport of cargo including whole organelles such as mitochondria between connected cells, intercellular nanotubes may create temporary syncytia, or ‘supercellularity’. Notably, syncytia are permanently created through cell-cell fusion, an essential event in a number of developmental and physiological processes such as in mammalian muscle and osteoclast function, and in the formation of the placental syncytiotrophoblast, a giant cell of ~12m^2^ surface area (Cox and Redman 2017). Intriguingly, cell fusion events have been linked to cellular senescence: the placental syncytiotrophoblast itself shows a number of markers of senescence (Cox and Redman 2017), while expression of fusogens such as endogenous retroviral ERVWE1 or measles virus induces cell fusion and induction of cell cycle arrest and senescence (Chuprin et al. 2013). Through induction of senescence, fused cells coordinate a protection strategy to prevent division of multinucleate cells, while maintaining cell viability through resisting apoptosis, which may be important within the placental syncytiotrophoblast throughout pregnancy (Chuprin et al. 2013) and in other physiological contexts such as in bone and muscle. It is interesting to speculate that through direct intercellular nanotubes, senescent cells could also relay danger signals to neighbouring cells and coordinate induction of paracrine senescence, just as cell fusion may induce senescence in syncytia. It will be of interest to assess the degree of electrical continuity between such connected senescent cells, as direct cytoplasmic bridges large enough for passage of mitochondria negate the need for other junctional structures such as gap junctions. Membrane potentials have been reported to be very low in senescence (Carroll et al. 2017), and this may have implications for the low propensity of senescent cells to undergo apoptosis. Direct cell-cell contacts leading to supercellularity are therefore of particular interest in senescence and in development of senomodifying therapies.

### Pathological relevance of direct intercellular contacts

While the data we present here suggest that a substantial proportion of senescent cells form direct cell-cell bridges which can mediate intercellular mitochondrial transfer, it is important to note that these data were acquired in 2-dimensional, *in vitro* cell culture of primary human skin and lung fibroblasts. While further work is undoubtedly required to probe the physiological relevance of these findings, previous *in vivo* work has shown that intercellular contacts can both rescue compromised cells and spread disease or infection, suggesting context-dependent function or exploitation by invading pathogens. Indeed, TNTs between tumour cells can play important roles in pathogenesis and invasion, for example through intercellular transfer of the P-glycoprotein drug efflux pump to propagate multi-drug resistance (Pasquier et al. 2012), together with intercellular mitochondrial transport for cellular rescue. TNTs have been implicated in the intercellular spread of pathogen such as HIV (Sowinski et al. 2008), influenza (Kumar et al. 2017) and herpes viral particles as well as prions (Gousset et al. 2009). Importantly in the context of age-related diseases, protein aggregates implicated in neurodegenerative diseases, including polyglutamine aggregates (Costanzo et al. 2013), α-synuclein (Wang et al. 2011, Abounit, Bousset, et al. 2016), and tau (Abounit, Wu, et al. 2016) have also been reported to be passed between cells through TNTs. The impact of intercellular tunnelling nanotubes is therefore likely to be determined by both the transferred cargo as well as the types and states of the connected cells.

### Possible roles of mitochondrial transfer through TNTs

Mitochondria are important signalling hubs for oxidative stress, apoptosis and senescence. While we note here transfer of mitochondria between cells, we did not address the functional status of the transferred mitochondria. Senescent cells are known to accumulate dysfunctional mitochondria, and previous reports suggest that healthy mitochondria may be preferentially transferred to stressed cells (Wang and Gerdes 2015). Measuring the O_2_ consumption and activity of individual ETC components in senescent and proliferating cells following co-culture would therefore be informative as to whether healthy or dysfunctional mitochondria are preferentially transferred. Another important line of enquiry is whether the nanotube structures that senescent cells make with NK cells (Wang and Gerdes 2015) also participate in intercellular mitochondrial transfer, as well as analysing the impact of direct interaction between senescent and proliferating cells, e.g. through analysis of proliferation capacity in co-cultures with direct contact compared with barrier trans-well formats. Moreover, it will be important to investigate the role of these intercellular connections in a physiologically relevant model, for example in 3D culture formats such as hydrogels of appropriate rigidity reflecting ‘young’ and ‘old’ tissue structures. Indeed, recently published data analysing TNT formation in mesenchymal stem cell (MSC) spheroids showed intercellular transfer of cytosolic dyes, and intriguingly, inclusion of low passage MSCs with high passage cells was seen to abrogate p16 expression in spheroids, dependent on TNT formation. Hence, in this context TNTs appear to rescue the proliferation potential of high passage cells, potentially overcoming senescence induction (Biran et al. 2015).

### Role of cell-cell communication in senescence

While senescent cells are known to accumulate in tissues with chronological age and within tumours, they also play important roles in development, tissue regeneration and in wound healing. Furthermore, the profile of cytokines, chemokines, matrix-remodelling enzymes and growth factors secreted in the SASP is highly heterogeneous between cell types and modes of senescence induction, perhaps underpinning the range of different paracrine responses to local induction of senescence. In these different physiological contexts, it is highly likely that senescent cells could exert a number of different impacts through direct intercellular bridges. Indeed, it is possible that within a tumour, intercellular bridges could participate in both rescuing cancer cells with dysfunctional mitochondria, but also in facilitating communication between senescent and immune cells to promote immune surveillance (Biran et al. 2015). It is also important to note that senescent cells have actually been shown to evade immune surveillance, for example by HLA-E upregulation (Pereira et al. 2019), thus TNTs may enable evasion of immune detection through shuttling of surface markers. Moreover, direct connections between senescent cells and proliferating neighbours may provide an additional means for spread of senescence – possibly even in the absence of secreted SASP factors that can induce bystander senescence. Therapies that focus on suppressing the SASP may therefore not be sufficient to ameliorate damaging aspects of senescent cell accumulation in cancer and ageing, and may additionally need to include approaches to disrupt direct cell-cell contacts through TNTs.

### Targeting intercellular tunnelling nanotubes in human disease

Proteins that promote actin polymerization and stabilization play an important role in facilitating direct cell-cell contact via membrane-bound bridges (Biran et al. 2015). Using co-culture assays and live cell imaging, we report here that not only do these bridges increase in frequency in senescence resulting from various stresses (replicative exhaustion, DNA damage or oncogene activation), but that they are regulated by mTOR and CDC42 signalling, and facilitate direct intercellular mitochondrial transfer. Importantly, Cdc42 may act downstream of constitutive mTOR activity in senescence.

While intercellular bridges may play important roles in physiological processes requiring cell-cell communication, such as aiding NK-mediated recognition of senescent cells (Chuprin et al. 2013), the pathological effects of cell-cell bridges may be amenable to therapeutic intervention. CDC42 signalling, implicated in the actin organization network, is reported to be elevated in senescent versus proliferating cells and to promote premature ageing (Wang et al. 2007), and as we show here, is necessary for TNT formation and/or stability. Actin cytoskeletal rearrangements, regulated by mTOR- and CD42-dependent pathways, are also likely to play a role in the enlarged, flattened morphology observed for senescent cells *in vitro* (Walters, Deneka-Hannemann, and Cox 2016), as well as enlargement *in vivo* (Biran et al. 2017), and our results show that senescent cells have higher numbers of TNTs than proliferating cells.

The importance of CDC42 and mTOR in senescence may therefore extend beyond regulating the SASP (Herranz et al. 2015, Laberge et al. 2015) and the biogenesis and secretion of exosomes (Zou et al. 2019), to promoting direct intercellular communication through formation of direct cell-cell communication channels. Consequently, our findings highlight mTOR signalling as critical in directing the non-cell-autonomous roles of cellular senescence, and further emphasise its validity as a therapeutic targe for senomodifying therapies, building on reports of beneficial effects of mTOR inhibitors on senescence phenotypes *in vitro* (Walters, Deneka-Hannemann, and Cox 2016), and longevity and health in vivo (Harrison et al. 2009, Urfer et al. 2017). It is tempting to speculate that the utility of mTOR inhibitors in a number of age-related diseases (reviewed in (Walters and Cox 2018)) including Alzheimer’s disease (Spilman et al. 2010) and immune senescence (Mannick et al. 2018), as well as their ability to improve ageing human skin structure (Chung et al. 2019), may be at least in part through actin cytoskeletal modulation that impacts on TNT formation.

Reducing TNT formation or stability through mTOR or CDC42 inhibition may be a useful therapeutic approach in cancer as well as in ageing, for example through blocking the transfer of healthy mitochondria from bystander cells to rescue dysfunctional tumour cells, or through blocking the spread of drug resistance channels. Indeed, neutralizing antibodies against the adhesion molecule ICAM-1 in T-cell acute lymphoblastic leukaemia were shown to block transfer of healthy mitochondria from MSCs, thereby causing an increase in chemotherapy-induced cancer cell death (Wang et al. 2018). This approach may also be important in preventing the spread of pathogens such as HIV or protein aggregates implicated in neurodegeneration: inhibiting nanotube formation with latrunculin B was shown to prevent the spread of α-synuclein through healthy astrocytes in a model of Parkinson’s disease propagation (Rostami et al. 2017).

Alternatively, it may be possible to exploit the beneficial properties of nanotubes, for example in treating diseases of organelle dysfunction, or by using intercellular mitochondrial transfer as a rescue strategy for compromised cells during stroke, when ischemic stress conditions may stimulate nanotube formation. In such circumstances, it may be possible to further promote nanotube formation by supplying a treatment that induces ROS, stabilizes microtubules or actin networks, or through inducing increased expression of trafficking adaptors in the mitochondria ‘donor’ cells. It may even be possible to exploit intercellular nanotubes as a drug delivery pathway in cancer therapy, as nanoparticles have been shown to be loaded into and move through TNTs (Kristl et al. 2013).

## Conclusions

In conclusion, we have demonstrated here a high prevalence of intercellular bridges (also called tunnelling nanotubes or TNTs) between senescent cells, that are membrane-bound and supported by an actin, and possibly also microtubule, cytoskeleton. Such bridges allow transfer of large cargo including mitochondria between cells. The dependence of such bridges on mTOR and CDC42 make them amenable to small molecule intervention, providing additional therapeutic strategies to modulate senescent cell behaviour in both age-related diseases and cancer.

## Data availability

Raw image files of all microscopy images shown, together with time lapse video of mitochondrial transfer between cells, and a Prism file of data for quantification of TNTs on drug treatment are available from the corresponding author upon email request.

## Acknowledgments

The authors gratefully acknowledge Mrs Anitha Nair for general cell culture support.

## Supplementary Materials

**Supplementary Figure:** Intercellular bridge formation is sensitive to CDC42 inhibition in a dose-dependent manner

## References

Abounit, S., L. Bousset, F. Loria, S. Zhu, F. de Chaumont, L. Pieri, J. C. Olivo-Marin, R. Melki, and C. Zurzolo. 2016. “Tunneling nanotubes spread fibrillar alpha-synuclein by intercellular trafficking of lysosomes.” EMBO J 35 (19):2120–2138. doi: 10.15252/embj.201593411.

Abounit, S., J. W. Wu, K. Duff, G. S. Victoria, and C. Zurzolo. 2016. “Tunneling nanotubes: A possible highway in the spreading of tau and other prion-like proteins in neurodegenerative diseases.” Prion 10 (5):344–351. doi: 10.1080/19336896.2016.1223003.

Arkwright, P. D., F. Luchetti, J. Tour, C. Roberts, R. Ayub, A. P. Morales, J. J. Rodriguez, A. Gilmore, B. Canonico, S. Papa, and M. D. Esposti. 2010. “Fas stimulation of T lymphocytes promotes rapid intercellular exchange of death signals via membrane nanotubes.” Cell Res 20 (1):72–88. doi: 10.1038/cr.2009.112.

Biran, A., M. Perelmutter, H. Gal, D. G. Burton, Y. Ovadya, E. Vadai, T. Geiger, and V. Krizhanovsky. 2015. “Senescent cells communicate via intercellular protein transfer.” Genes Dev 29 (8):791–802. doi: 10.1101/gad.259341.115.

Biran, A., L. Zada, P. Abou Karam, E. Vadai, L. Roitman, Y. Ovadya, Z. Porat, and V. Krizhanovsky. 2017. “Quantitative identification of senescent cells in aging and disease.” Aging Cell 16 (4):661–671. doi: 10.1111/acel.12592.

Carroll, B., G. Nelson, Y. Rabanal-Ruiz, O. Kucheryavenko, N. A. Dunhill-Turner, C. C. Chesterman, Q. Zahari, T. Zhang, S. E. Conduit, C. A. Mitchell, O. D. K. Maddocks, P. Lovat, T. von Zglinicki, and V. I. Korolchuk. 2017. “Persistent mTORC1 signaling in cell senescence results from defects in amino acid and growth factor sensing.” J Cell Biol 216 (7):1949–1957. doi: 10.1083/jcb.201610113.

Chinnery, H. R., E. Pearlman, and P. G. McMenamin. 2008. “Cutting edge: Membrane nanotubes in vivo: a feature of MHC class II+ cells in the mouse cornea.” J Immunol 180 (9):5779–83. doi: 10.4049/jimmunol.180.9.5779.

Chung, Christina Lee, Ibiyonu Lawrence, Melissa Hoffman, Dareen Elgindi, Kumar Nadhan, Manali Potnis, Annie Jin, Catlin Sershon, Rhonda Binnebose, Antonello Lorenzini, and Christian Sell. 2019. “Topical rapamycin reduces markers of senescence and aging in human skin: an exploratory, prospective, randomized trial.” GeroScience 41 (6):861–869. doi: 10.1007/s11357-019-00113-y.

Chuprin, A., H. Gal, T. Biron-Shental, A. Biran, A. Amiel, S. Rozenblatt, and V. Krizhanovsky. 2013. “Cell fusion induced by ERVWE1 or measles virus causes cellular senescence.” Genes Dev 27 (21):2356–66. doi: 10.1101/gad.227512.113.

Correia-Melo, C., F. D. Marques, R. Anderson, G. Hewitt, R. Hewitt, J. Cole, B. M. Carroll, S. Miwa, J. Birch, A. Merz, M. D. Rushton, M. Charles, D. Jurk, S. W. Tait, R. Czapiewski, L. Greaves, G. Nelson, Y. M. Bohlooly, S. Rodriguez-Cuenca, A. Vidal-Puig, D. Mann, G. Saretzki, G. Quarato, D. R. Green, P. D. Adams, T. von Zglinicki, V. I. Korolchuk, and J. F. Passos. 2016. “Mitochondria are required for pro-ageing features of the senescent phenotype.” EMBO J 35 (7):724–42. doi: 10.15252/embj.201592862.

Costanzo, M., S. Abounit, L. Marzo, A. Danckaert, Z. Chamoun, P. Roux, and C. Zurzolo. 2013. “Transfer of polyglutamine aggregates in neuronal cells occurs in tunneling nanotubes.” J Cell Sci 126 (Pt 16):3678–85. doi: 10.1242/jcs.126086.

Cox, L. S., and C. Redman. 2017. “The role of cellular senescence in ageing of the placenta.” Placenta 52:139–145. doi: 10.1016/j.placenta.2017.01.116.

de Magalhães, João Pedro, and João F. Passos. 2018. “Stress, cell senescence and organismal ageing.” Mechanisms of Ageing and Development 170:2–9. doi: https://doi.org/10.1016/j.mad.2017.07.001.

Domhan, S., L. Ma, A. Tai, Z. Anaya, A. Beheshti, M. Zeier, L. Hlatky, and A. Abdollahi. 2011. “Intercellular communication by exchange of cytoplasmic material via tunneling nano-tube like structures in primary human renal epithelial cells.” PLoS One 6 (6):e21283. doi: 10.1371/journal.pone.0021283.

Dupont, M., S. Souriant, G. Lugo-Villarino, I. Maridonneau-Parini, and C. Verollet. 2018. “Tunneling Nanotubes: Intimate Communication between Myeloid Cells.” Front Immunol 9:43. doi: 10.3389/fimmu.2018.00043.

Gonzalez-Meljem, J. M., J. R. Apps, H. C. Fraser, and J. P. Martinez-Barbera. 2018. “Paracrine roles of cellular senescence in promoting tumourigenesis.” Br J Cancer 118 (10):1283–1288. doi: 10.1038/s41416-018-0066-1.

Gousset, K., E. Schiff, C. Langevin, Z. Marijanovic, A. Caputo, D. T. Browman, N. Chenouard, F. de Chaumont, A. Martino, J. Enninga, J. C. Olivo-Marin, D. Mannel, and C. Zurzolo. 2009. “Prions hijack tunnelling nanotubes for intercellular spread.” Nat Cell Biol 11 (3):328–36. doi: 10.1038/ncb1841.

Hanna, S. J., K. McCoy-Simandle, V. Miskolci, P. Guo, M. Cammer, L. Hodgson, and D. Cox. 2017. “The Role of Rho-GTPases and actin polymerization during Macrophage Tunneling Nanotube Biogenesis.” Sci Rep 7 (1):8547. doi: 10.1038/s41598-017-08950-7.

Harrison, David E., Randy Strong, Zelton Dave Sharp, James F. Nelson, Clinton M. Astle, Kevin Flurkey, Nancy L. Nadon, J. Erby Wilkinson, Krystyna Frenkel, Christy S. Carter, Marco Pahor, Martin A. Javors, Elizabeth Fernandez, and Richard A. Miller. 2009. “Rapamycin fed late in life extends lifespan in genetically heterogeneous mice.” Nature 460 (7253):392–395. doi: 10.1038/nature08221.

Herranz, N., S. Gallage, M. Mellone, T. Wuestefeld, S. Klotz, C. J. Hanley, S. Raguz, J. C. Acosta, A. J. Innes, A. Banito, A. Georgilis, A. Montoya, K. Wolter, G. Dharmalingam, P. Faull, T. Carroll, J. P. Martinez-Barbera, P. Cutillas, F. Reisinger, M. Heikenwalder, R. A. Miller, D. Withers, L. Zender, G. J. Thomas, and J. Gil. 2015. “mTOR regulates MAPKAPK2 translation to control the senescence-associated secretory phenotype.” Nat Cell Biol 17 (9):1205–17. doi: 10.1038/ncb3225.

Korolchuk, Viktor I., Satomi Miwa, Bernadette Carroll, and Thomas von Zglinicki. 2017. “Mitochondria in Cell Senescence: Is Mitophagy the Weakest Link?” EBioMedicine 21:7–13. doi: 10.1016/j.ebiom.2017.03.020.

Koyanagi, M., R. P. Brandes, J. Haendeler, A. M. Zeiher, and S. Dimmeler. 2005. “Cell-to-cell connection of endothelial progenitor cells with cardiac myocytes by nanotubes: a novel mechanism for cell fate changes?” Circ Res 96 (10):1039–41. doi: 10.1161/01.RES.0000168650.23479.0c.

Kristl, J., K. T. Plajnsek, M. E. Kreft, B. Jankovic, and P. Kocbek. 2013. “Intracellular trafficking of solid lipid nanoparticles and their distribution between cells through tunneling nanotubes.” Eur J Pharm Sci 50 (1):139–48. doi: 10.1016/j.ejps.2013.04.013.

Kumar, A., J. H. Kim, P. Ranjan, M. G. Metcalfe, W. Cao, M. Mishina, S. Gangappa, Z. Guo, E. S. Boyden, S. Zaki, I. York, A. Garcia-Sastre, M. Shaw, and S. Sambhara. 2017. “Influenza virus exploits tunneling nanotubes for cell-to-cell spread.” Sci Rep 7:40360. doi: 10.1038/srep40360.

Laberge, R. M., Y. Sun, A. V. Orjalo, C. K. Patil, A. Freund, L. Zhou, S. C. Curran, A. R. Davalos, K. A. Wilson-Edell, S. Liu, C. Limbad, M. Demaria, P. Li, G. B. Hubbard, Y. Ikeno, M. Javors, P. Y. Desprez, C. C. Benz, P. Kapahi, P. S. Nelson, and J. Campisi. 2015. “MTOR regulates the pro-tumorigenic senescence-associated secretory phenotype by promoting IL1A translation.” Nat Cell Biol 17 (8):1049–61. doi: 10.1038/ncb3195.

Lou, E., S. Fujisawa, A. Morozov, A. Barlas, Y. Romin, Y. Dogan, S. Gholami, A. L. Moreira, K. Manova-Todorova, and M. A. Moore. 2012. “Tunneling nanotubes provide a unique conduit for intercellular transfer of cellular contents in human malignant pleural mesothelioma.” PLoS One 7 (3):e33093. doi: 10.1371/journal.pone.0033093.

Mannick, J. B., M. Morris, H. P. Hockey, G. Roma, M. Beibel, K. Kulmatycki, M. Watkins, T. Shavlakadze, W. Zhou, D. Quinn, D. J. Glass, and L. B. Klickstein. 2018. “TORC1 inhibition enhances immune function and reduces infections in the elderly.” Sci Transl Med 10 (449). doi: 10.1126/scitranslmed.aaq1564.

Marzo, L., K. Gousset, and C. Zurzolo. 2012. “Multifaceted roles of tunneling nanotubes in intercellular communication.” Front Physiol 3:72. doi: 10.3389/fphys.2012.00072.

Nelson, G., J. Wordsworth, C. Wang, D. Jurk, C. Lawless, C. Martin-Ruiz, and T. von Zglinicki. 2012. “A senescent cell bystander effect: senescence-induced senescence.” Aging Cell 11 (2):345–9. doi: 10.1111/j.1474-9726.2012.00795.x.

Ohno, H., K. Hase, and S. Kimura. 2010. “M-Sec: Emerging secrets of tunneling nanotube formation.” Commun Integr Biol 3 (3):231–3. doi: 10.4161/cib.3.3.11242.

Onfelt, B., S. Nedvetzki, R. K. Benninger, M. A. Purbhoo, S. Sowinski, A. N. Hume, M. C. Seabra, M. A. Neil, P. M. French, and D. M. Davis. 2006. “Structurally distinct membrane nanotubes between human macrophages support long-distance vesicular traffic or surfing of bacteria.” J Immunol 177 (12):8476–83. doi: 10.4049/jimmunol.177.12.8476.

Ovadya, Y., T. Landsberger, H. Leins, E. Vadai, H. Gal, A. Biran, R. Yosef, A. Sagiv, A. Agrawal, A. Shapira, J. Windheim, M. Tsoory, R. Schirmbeck, I. Amit, H. Geiger, and V. Krizhanovsky. 2018. “Impaired immune surveillance accelerates accumulation of senescent cells and aging.” Nat Commun 9 (1):5435. doi: 10.1038/s41467-018-07825-3.

Pasquier, J., L. Galas, C. Boulange-Lecomte, D. Rioult, F. Bultelle, P. Magal, G. Webb, and F. Le Foll. 2012. “Different modalities of intercellular membrane exchanges mediate cell-to-cell p-glycoprotein transfers in MCF-7 breast cancer cells.” J Biol Chem 287 (10):7374–87. doi: 10.1074/jbc.M111.312157.

Pereira, B. I., O. P. Devine, M. Vukmanovic-Stejic, E. S. Chambers, P. Subramanian, N. Patel, A. Virasami, N. J. Sebire, V. Kinsler, A. Valdovinos, C. J. LeSaux, J. F. Passos, A. Antoniou, M. H. A. Rustin, J. Campisi, and A. N. Akbar. 2019. “Senescent cells evade immune clearance via HLA-E-mediated NK and CD8(+) T cell inhibition.” Nat Commun 10 (1):2387. doi: 10.1038/s41467-019-10335-5.

Rostami, J., S. Holmqvist, V. Lindstrom, J. Sigvardson, G. T. Westermark, M. Ingelsson, J. Bergstrom, L. Roybon, and A. Erlandsson. 2017. “Human Astrocytes Transfer Aggregated Alpha-Synuclein via Tunneling Nanotubes.” J Neurosci 37 (49):11835–11853. doi: 10.1523/JNEUROSCI.0983-17.2017.

Rustom, A., R. Saffrich, I. Markovic, P. Walther, and H. H. Gerdes. 2004. “Nanotubular highways for intercellular organelle transport.” Science 303 (5660):1007–10. doi: 10.1126/science.1093133.

Sowinski, S., C. Jolly, O. Berninghausen, M. A. Purbhoo, A. Chauveau, K. Kohler, S. Oddos, P. Eissmann, F. M. Brodsky, C. Hopkins, B. Onfelt, Q. Sattentau, and D. M. Davis. 2008. “Membrane nanotubes physically connect T cells over long distances presenting a novel route for HIV-1 transmission.” Nat Cell Biol 10 (2):211–9. doi: 10.1038/ncb1682.

Spilman, P., N. Podlutskaya, M. J. Hart, J. Debnath, O. Gorostiza, D. Bredesen, A. Richardson, R. Strong, and V. Galvan. 2010. “Inhibition of mTOR by rapamycin abolishes cognitive deficits and reduces amyloid-beta levels in a mouse model of Alzheimer’s disease.” PLoS One 5 (4):e9979. doi: 10.1371/journal.pone.0009979.

Takahashi, A., A. Kukita, Y. J. Li, J. Q. Zhang, H. Nomiyama, T. Yamaza, Y. Ayukawa, K. Koyano, and T. Kukita. 2013. “Tunneling nanotube formation is essential for the regulation of osteoclastogenesis.” J Cell Biochem 114 (6):1238–47. doi: 10.1002/jcb.24433.

Uphoff, C. C., and H. G. Drexler. 2002. “Comparative PCR analysis for detection of mycoplasma infections in continuous cell lines.” In Vitro Cell Dev Biol Anim 38 (2):79–85. doi: 10.1290/1071-2690(2002)038<0079:CPAFDO>2.0.CO;2.

Uphoff, C. C., and H. G. Drexler. 2004. “Detecting Mycoplasma contamination in cell cultures by polymerase chain reaction.” Methods Mol Med 88:319–26. doi: 10.1385/1-59259-406-9:319.

Urfer, Silvan R., Tammi L. Kaeberlein, Susan Mailheau, Philip J. Bergman, Kate E. Creevy, Daniel E. L. Promislow, and Matt Kaeberlein. 2017. “A randomized controlled trial to establish effects of short-term rapamycin treatment in 24 middle-aged companion dogs.” GeroScience 39 (2):117–127. doi: 10.1007/s11357-017-9972-z.

Veranic, P., M. Lokar, G. J. Schutz, J. Weghuber, S. Wieser, H. Hagerstrand, V. Kralj-Iglic, and A. Iglic. 2008. “Different types of cell-to-cell connections mediated by nanotubular structures.” Biophys J 95 (9):4416–25. doi: 10.1529/biophysj.108.131375.

Walters, H. E., and L. S. Cox. 2018. “mTORC Inhibitors as Broad-Spectrum Therapeutics for Age-Related Diseases.” Int J Mol Sci 19 (8):2325. doi: 10.3390/ijms19082325.

Walters, H. E., S. Deneka-Hannemann, and L. S. Cox. 2016. “Reversal of phenotypes of cellular senescence by pan-mTOR inhibition.” Aging (Albany NY) 8 (2):231–44. doi: 10.18632/aging.100872.

Wang, J., X. Liu, Y. Qiu, Y. Shi, J. Cai, B. Wang, X. Wei, Q. Ke, X. Sui, Y. Wang, Y. Huang, H. Li, T. Wang, R. Lin, Q. Liu, and A. P. Xiang. 2018. “Cell adhesion-mediated mitochondria transfer contributes to mesenchymal stem cell-induced chemoresistance on T cell acute lymphoblastic leukemia cells.” J Hematol Oncol 11 (1):11. doi: 10.1186/s13045-018-0554-z.

Wang, Lei, Linda Yang, Marcella Debidda, David Witte, and Yi Zheng. 2007. “Cdc42 GTPase-activating protein deficiency promotes genomic instability and premature aging-like phenotypes.” Proceedings of the National Academy of Sciences 104 (4):1248. doi: 10.1073/pnas.0609149104.

Wang, X., and H. H. Gerdes. 2015. “Transfer of mitochondria via tunneling nanotubes rescues apoptotic PC12 cells.” Cell Death Differ 22 (7):1181–91. doi: 10.1038/cdd.2014.211.

Wang, Y., J. Cui, X. Sun, and Y. Zhang. 2011. “Tunneling-nanotube development in astrocytes depends on p53 activation.” Cell Death Differ 18 (4):732–42. doi: 10.1038/cdd.2010.147.

Zou, W., M. Lai, Y. Zhang, L. Zheng, Z. Xing, T. Li, Z. Zou, Q. Song, X. Zhao, L. Xia, J. Yang, A. Liu, H. Zhang, Z. K. Cui, Y. Jiang, and X. Bai. 2019. “Exosome Release Is Regulated by mTORC1.” Adv Sci (Weinh) 6 (3):1801313. doi: 10.1002/advs.201801313.

