## Supplementary figure for "Intercellular transfer of mitochondria between senescent cells through cytoskeleton-supported intercellular bridges requires mTOR and Cdc42 signalling"

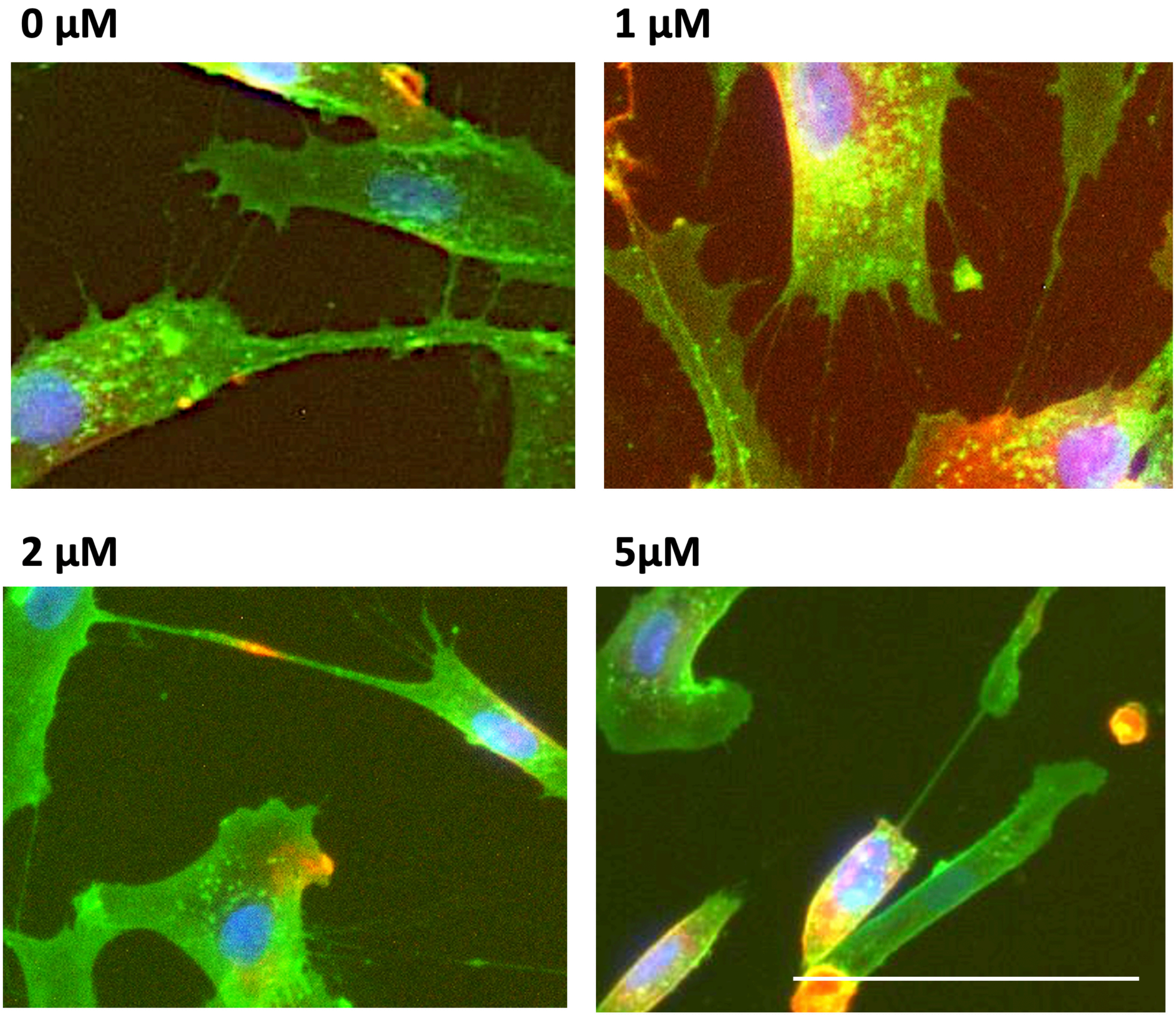


**Supplementary Figure: Intercellular bridge formation is sensitive to CDC42 inhibition in a dose-dependent manner.** HF043 fibroblasts at CPD 71 were incubated with 0, 1, 2 and 5 µm CASIN, an inhibitor of CDC42 for 24h then stained with FITC-WGA (membranes), rhodamine phalloidin (actin) and NucBlue Live for DNA. Scale bar 100 µm.
